# The Multiple Approaches for Drug-Drug Interaction Extraction using Machine learning and transformer based Model

**DOI:** 10.1101/2025.10.25.684550

**Authors:** Gurpreet Singh, Sundareswari Thiyagarajan

## Abstract

This research paper investigates a machine learning based approach for Drug-Drug Interaction (DDI) extraction for determining the side effects of multi drugs when prescribed simultaneously. In our proposed model we used TAC 2017 Dataset, which has Adverse Drug Reactions (ADRs) data for the classification ofdrug-drug interaction. TAC 2017 Dataset has various types of information which are related to drugs and their interactions. Our method uses Term Frequency-Inverse Document Frequency (TF-IDF) to transform the textual descriptions ofside effects for DDI into numerical feature vectors, followed by a Random Forest Classifier, Gradient Boosting, BioBERT, Support Vector Machine (SVM) Algorithms to predict the potential interactions between drug pairs. One of the key strength of the Random Forest approach is its ability to provide feature importance scores, which allows us to interpret which side effects are most influential in predicting drug interactions. The key advantage of Gradient Boosting is its high predictive performance combined with interpretability. It is able to handle complex, structured data efficiently. Additionally, the model’s decisions are more transparent, which is necessary in the biomedical domain. The advantageof SVM is its ability to handle high-dimensional data, capture complex non-linear interactions using kernel functions, and generalize with datasets, making it robust to over-fitting. BioBERT (Bidirectional Encoder Representations from Transformers for Biomedical Text Mining) is advantageous for DDI prediction due to its biomedical domain knowledge, contextual understanding of complex drug-related texts.These method captures the relevance and importance of each side effect of multi drugs and also generate pairs of drugs from the dataset. Our model demonstrates competitive performance in DDI prediction, which highlights the utility of text-based feature extraction combined with an interpretable ensemble learning model.

## I. Introduction

Drug-Drug Interaction (DDI) refers to effect of consuming two or more drugs simultaneously. There is a study on DDI pairs in vivo and in vitro research. Laboratory and practical research can give more accurate results but on the other hand, it was expensive and also has limited resources [12]. Drug-drug interactions (DDI) plays a vital role in the development of new drugs and also for (PhV) pharmacovigilance which contributes 6 - 30% of all Adverse Drug Events [13]. There are many methods proposed for DDI extraction that are classified into two categories such as rule-based [16] and machine learning-based methods [17], [19]. In comparison between better performance and portability Machine learning models outperform the rule-based models [18]. BERT [43] is a recently proposed pre-trained transformer based model. Due to its multi-layer bidirectional transformer [44] structure, BERT can process the contextual information of sentences into the word vector both from forward and backwards. Classification-based methods formulate DDI prediction as a binary classification task which is represented drug-drug pairs as feature vectors from their constructed pharma cointeraction network [45]. Then they defined the presence or absence of interactions as labels and finally built the logistic regression models for predictions.

In this study, we propose a method using Term Frequency-Inverse Document Frequency (TF-IDF) vectorization combined with Random Forest Classifier, SVM, Gradient Boosting, while BioBERT combined with neural network to predict DDIs. By training the Random Forest model on the feature vectors, we focus to predict whether a given pair of drugs will interact or not. A binary classification label is assigned based on whether the drugs share common side effects or not, indicating a potential interaction. Each model offers distinct advantages,traditional Machine Learning methods are efficient for structured data, while BioBERT leverages deep learning to analyze the unstructured text.

## II. Literature Review

From between the drug pairs identifying the drug entities and their interaction is one of the challenging tasks for Drug-Drug Interaction (DDI) Extraction [20]. For the Drug Discovery Computer-Aided Drug-Drug Interaction (DDI) Extraction plays a vital role, because of its process is too expensive and also an time-consuming. So that, in the drug development cycle Machine Learning based approaches can reduce the laborious task. Here the author proposed an approach namely TP-DDI (TP-DDI: Transformer-based pipeline for the extraction of Drug-Drug Interactions) using an Pre-trained weights called BioBERT to improve both drug name entity recognition and DDI extraction task on the DDI Extraction 2013 dataset. In [14], Drug-Drug Interaction (DDI) Extraction is one of the Natural Language Processing task. The author proposed an CNN (Convolutional Neural Network) based model DDI Extraction using DDI 2013 Corpus dataset, their proposed model achieved an F-score of 69.75%.In [21], Preventing adverse effects from drug combinations can be achieve by detecting drug-drug interactions (DDI). The author proposed an Recursive Neural Network to improve the performance of DDI extraction using DDI 2013 dataset. Their proposed model comprises a position feature, a subtree containment feature, and an ensemble method to improve the performance of DDI extraction. They compared with the state-of-the-art models, the DDI detection and type classifiers of their model performed 4.4% and 2.8% better than the existing model. In [22] DDI extraction means extracting information from biomedical text it is a promising approach it gives the better insights to know about the effect of one drug with another drug. However the available of large data makes it more complicated to understand Drug-Drug Interaction and its side effects. Here the author proposed an text mining method for extracting automatically DDI information from text. They combined feature embedding with deep learning methods, and also they achieved improving results in learning the text features by using an LSTM (Long-short term memory) neural network. In [23] Here the author proposes a novel solution to DDI Extraction by using Relation BioBERT (R-BioBERT) to detect and classify Drug-Drug Interaction with the Bidirectional Long Short-Term Memory (BLSTM) for improving the accuracy of predictions for the Drug-Drug Interaction. Their proposed model also can able to identifies the specific type of interaction between the two drug interaction. Their proposed model shows that the use of BLSTM yields higher F-scores compared to their baseline model using the three well-known dataset such as SemEval 2013, TAC 2018, and TAC 2019. In [24], the healthcare system, medical errors is one of the growing concerns because of its adverse events. Drug-Drug Interaction can able to prevent adverse events, by extracting Drug-Drug Interactions from drug labels into a machine-processable form, its an effective step towards an drug safety information in the health care system. They proposed an graph convolutions (GCs) with a novel attention-based gating mechanism (GCA) for predicting task using TAC 2018 DDI Extraction corpus.In [25], In this paper the authors proposed three deep learning models for the classification and identification method for Drug-Drug Interaction such as CNN (Convolutional Neural Network), BiLSTM (Bi-directional Long-Short term Memory), and BiLSTM with attention. They also introduce three transformer-based models namely BERT, RoBERTa, and ELECTRA. To improve the DDI extraction they proposed an ensemble approach by combining these transformer models using DDI Extraction 2013 dataset. And also their attention-based model gives better yield compared to their existing model.

## III. Research Method and Measures

**Fig. 1.**
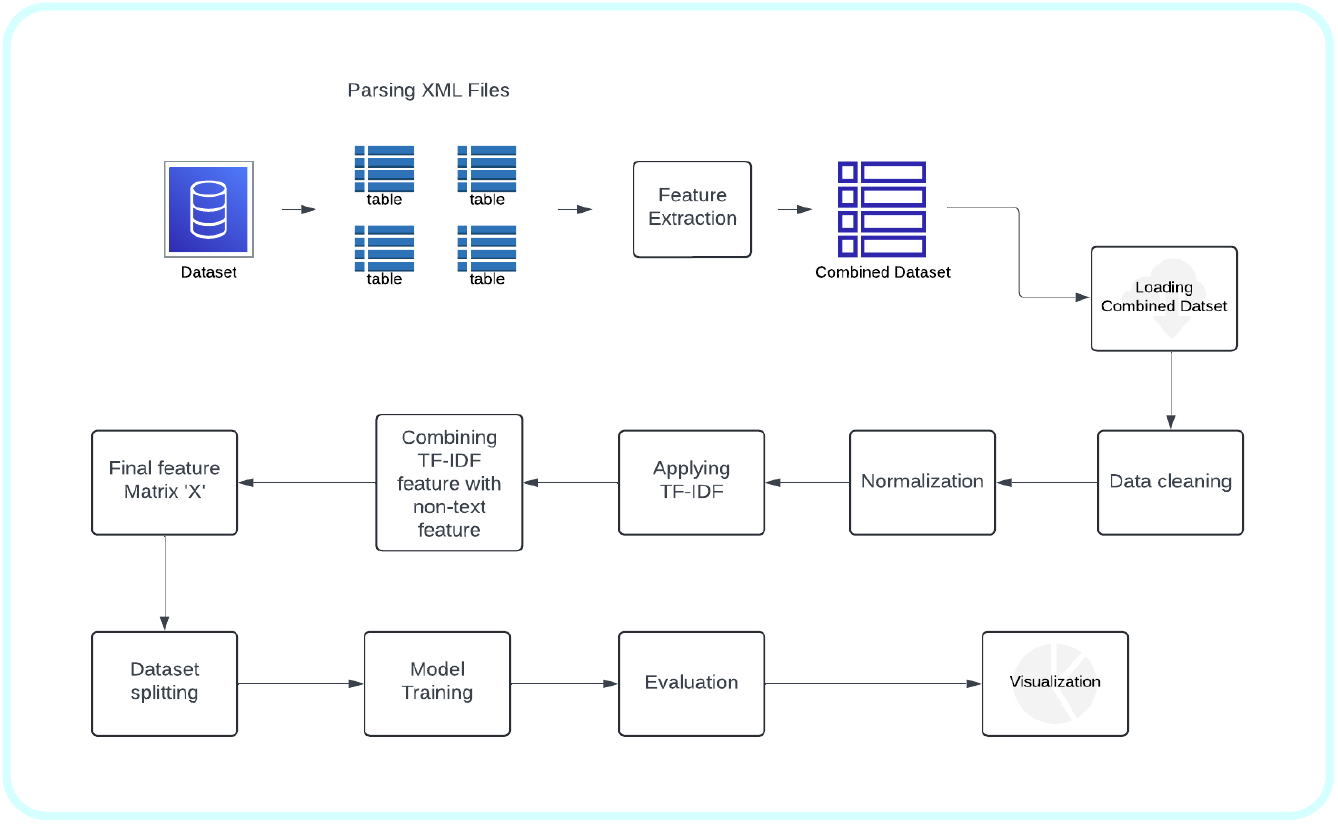
Research Model

### 3.1 Dataset Description

This TAC 2017 Adverse Drug Reaction dataset is one of the part of the Text Analysis Conference (TAC) Adverse Drug Reaction Extraction from Drug Labels(Drug prescription) which describe how to use, who should not use, who should use, the reaction from the particular medicine and some specific safety concerns. This dataset is primarily focused on extracting and normalizing adverse drug reactions (ADRs) from structured product labels (SPLs) of the US Food and Drug Administration (FDA) drugs. This dataset includes annotations for ADR(Adverse Drug Reaction), the relation between ADR such as hypothetical, effect, and negated, and their reactions. These Adverse Drug Reactions are then normalized according to the Medical Dictionary in terms of MedDRA terminology, to provide consistency and facilitation for further analysis. These datasets are composed of SPLs (structured product labels), which are one of the official documents, that describe the drug safety, effect, and usage of drugs.

For our proposed model we used TAC 2017 Dataset for the Multi-class classification of drug-drug interaction, TAC 2017 Dataset has various types of information which are related to drugs and their interactions. The dataset consists of Textual Features and Integer Features. Textual Features such as Drug, Arg1, Arg2, Reaction_Str, Normalization_id, Meddra_PT, Meddra_LLT, which contains the drug information,their relation and their reactions. Integers Features such as Meddra_PT_ID, Meddra_LLT_ID these are numerical codes along with medical terms, it is basically used for categorization and analysis.

**Fig. 2.**
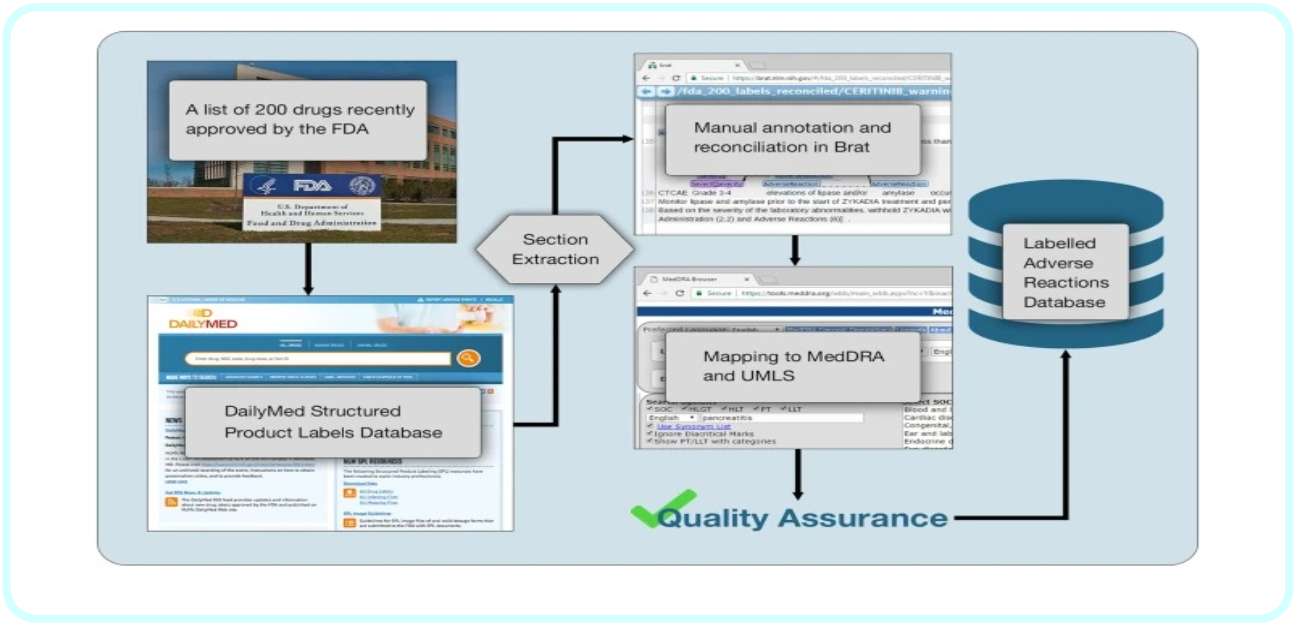
Representation of ADR Database [41]

### 3.2 Data Preprocessing

The textual features which was processed by TF-IDF were combined with the integer features into a single feature matrix. By doing this the model can able to learn the data from both textual and numerical data. We combined the dataset which comprises various Drugs and their relation in a separate folder. In each and every folder, it contains details about each drug’s properties, such as its name, relation, and reaction or interaction properties, which contains information about the drugs interactions, including whether the interaction is hypothetical, effectual, or negated with annotated fields along with Meddra Codes, The MedDRA (Medical Dictionary for Regulatory Activities) codes which is refers to standardized medical terms for adverse events and the medical conditions, which are used to describe the interactions between the drugs.

**Fig. 3.**
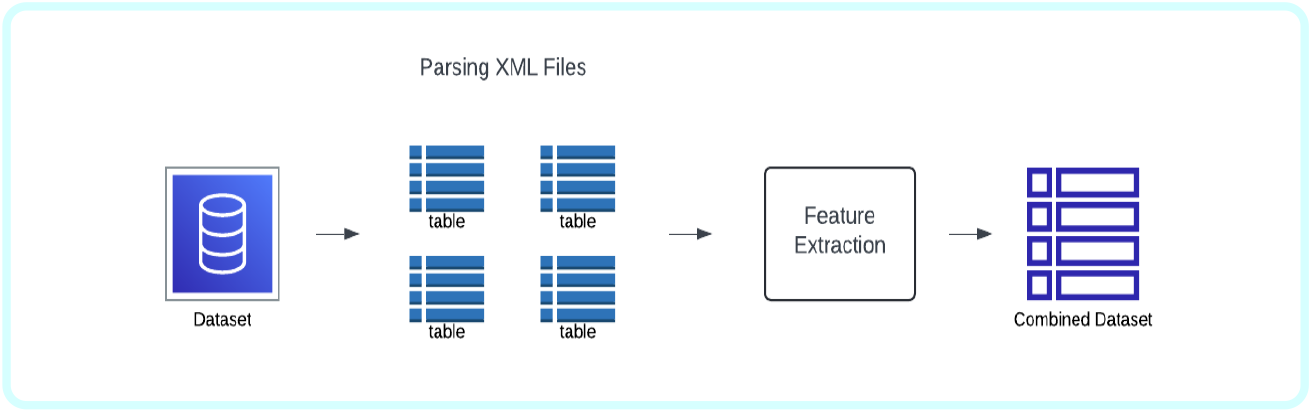
Visual representation of Data preprocessing

### 3.4 Combination of data

Textual columns in this dataset were combined into a single column. This combined column comprises the drug and its side effects. Empty strings were placed in place of missing values to ensure valid text representation for each and every data.

### 3.5 Feature Extraction

These text data were converted into numerical features applying TF-IDF (Term Frequency-Inverse Document Frequency). By using this technique it converts the text data in the form of a matrix of TF-IDF vectorization, these matrix represent the importance of each word present in the data. To control the complexity of model computational and its dimension we limited the maximum number of features into 2000 and also we removed English stop word to improve the focus on the main or meaningful word.

For our model, we used TF-IDF (Term Frequency-Inverse Document Frequency), which is one of the methods in Natural Language Processing (NLP) used for converting textual data into numerical features. TF means Term Frequency it is used to measure the frequent words in files. IDF means Inverse Document Frequency which is used to measure important words in the entire dataset by decrementing the weight of normal words in files. It will give the result in the matrix form where each row denotes the files about the drug-drug interaction while each column denotes a word, which is weighted by its importance in files. TF-IDF stands for Term Frequency-Inverse Document Frequency. It is vastly used in information retrieval, NLP, text classification, keyword extraction [2-3].

Term Frequency (TF) is used to count how frequently a text comes in a document. Term is represented as ‘t’ and Document as ‘d’.

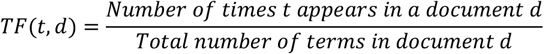

IDF measures the importance of a particular term in the entire document. It is used in reducing the common terms (“are”, “is”,”It”, etc.) which occurs frequently. |D| is the Total number of documents in the corpus. Number of documents containing term t refers to the count of documents in which the term t appears.

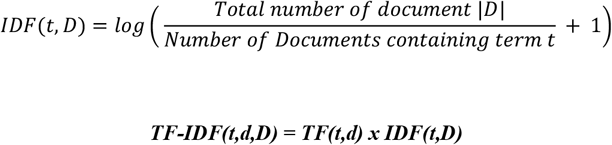

TF-IDF Score is calculated by multiplying the TF and IDF. High TF-IDF Score refers to the good keyword for the document and low TF-IDF Score refers to the less informative term.

### 3.6 Normalization

After TF-IDF vectorization these features were normalized using L2 normalization. By using normalization it helps in balancing the instances of unique features during the model training.

### 3.7 Data Splitting

By using an 80/20 split we split the dataset into training and test sets, training set for model training, while the testing set for evaluating the model performance.

## IV. Evaluation Metrices

True positive (TP), False Positive (FP), True Negative (TN), False Negative (FN) are the important factors in Evaluation of Accuracy, Precision, Recall, F1-score.

True Positive (TP) refers to **t**he model correctly identifies the interaction from the dataset. False Positive (FP) refers to **t**he model incorrectly identifies no interaction from the dataset. True Negative (TN) refers to the model correctly identifies no interaction from the dataset. False Negative (FN) refers to **t**he model incorrectly identifies the interaction from the dataset.

### 4.1 Accuracy

It shows the overall performance of our model by correctly classifying instances.

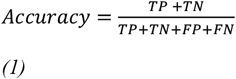

### 4.2 Precision

Precision is also known as Positive Predictive Value. It is the ratio of correctly predicted positive value to the total predicted positive value. It is effective only the cost of false positive value is high.

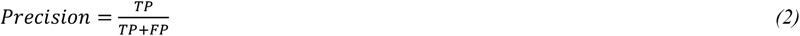

### 4.3 Recall

Recall is also known as True Positive Rate. Recall refers to the ratio of correctly predicted positive cases to the total number of positive instances

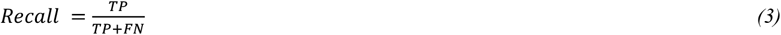

### 4.4 F1-Score

It shows the metrics between precision and recall. It is most useful when we are dealing with imbalanced datasets for training the model for multi-class classification.

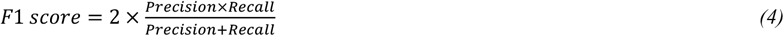

### 4.5 ROC along with AUC

AUC (Area Under the Curve) of the ROC (Receiver Operating Characteristic) curve, which measures the model’s ability to differentiate between the classes across all possible classification thresholds.t shows the difference between the True Positive Rate and False Positive Rate for the threshold values.

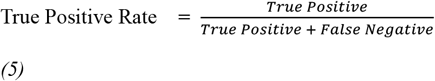

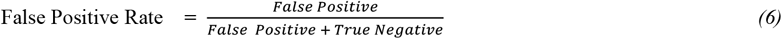

## V. Model Training and Evaluation

### 5.1 Random Forest Algorithm and its Performance

For our proposed model we used Random Forest Classifier to perform the classification tasks. Random forest classifier is an ensemble learning method, it combines multiple decision trees to improve performance and reduce overfitting. Using Randomized Search CV we perform tuning for the hyperparameters for model training. Random forest is an ensemble of decision trees that has multiple processes like bagging, random feature selection [5-8]. It is combination of the decision trees for the purpose of increasing the accuracy, reducing the over-fitting. It does not have a structure like single closed-form formula. The final classification or prediction is made based on the majority of the voting among all the decision trees. Most of the decision is based on the bagging and random feature selection, which will helps in reducing the variance and generalization.

#### 5.1.1 Bagging (Bootstrap Sampling)

Bagging is done by creating multiple datasets by randomly sampling the original with replacement. Each tree in Random Forest Algorithm is trained on a different subset of the data.

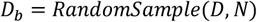

Where,

N: Total number of data points

*D*_*b*_: Bootstraped sample for tree b

*D*: The original dataset

#### 5.1.2 Random Feature Selection

For every split in the decision tree, a random subset is considered for performing best split. ‘p’ represents the total number of features. ‘m’ represents the number of features selected at each split.

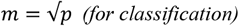

Where,

p - total number of features

m - number of features selected at each split.

#### 5.1.3 Ensemble Voting for Prediction (Majority Voting)

The final prediction is made by aggregation prediction from all the decision trees.

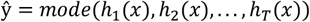

Where,

*ŷ* - Overall Prediction

*h*_*i*_(*x*) - Class prediction

Some of the key hyper-parameters tuning include the number of estimators, maximum depth, minimum samples split, and minimum samples leaf. After model training, it visualizes the performance evaluation metrics.

**Fig. 4.**
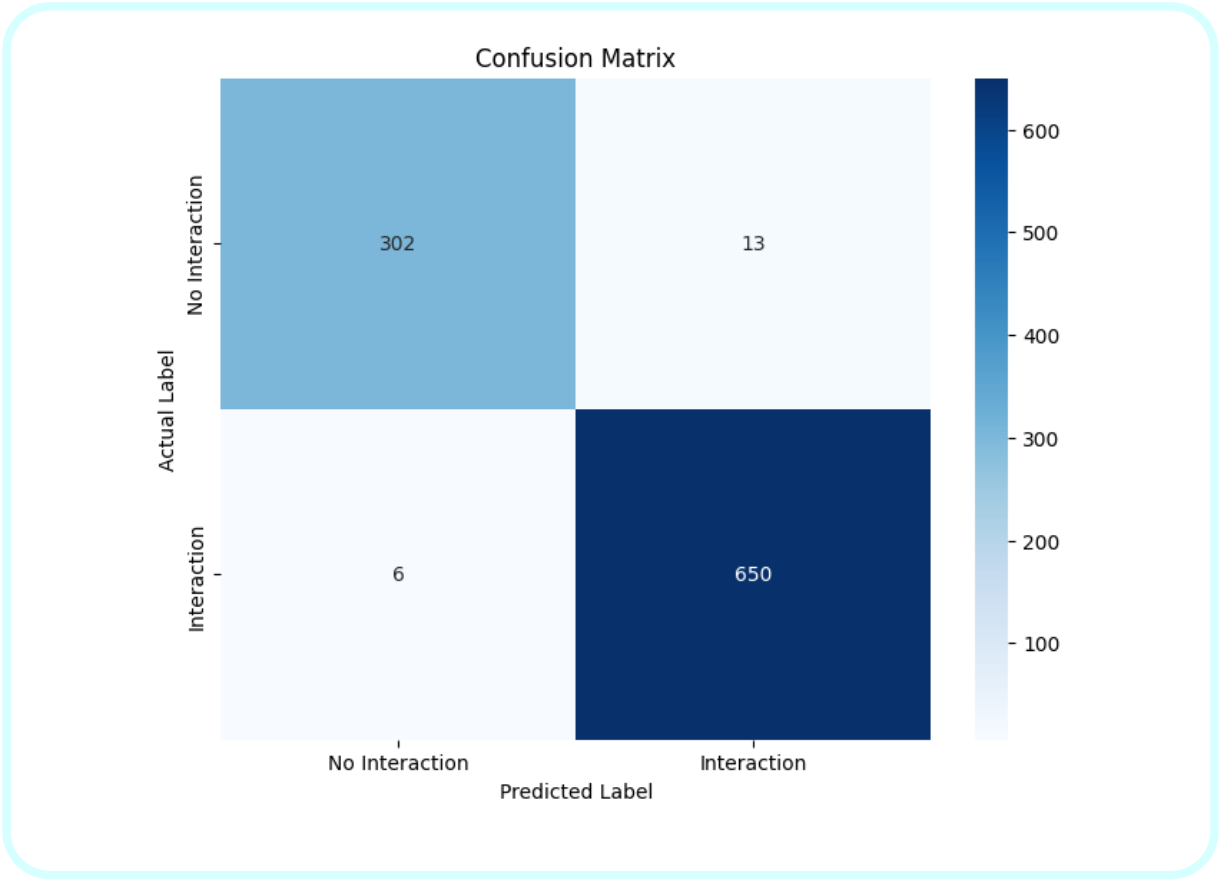
Confusion matrix for Random Forest

True Positive (TP) = 650, False Positive (FP) = 13, True Negative(TN) = 302, False Negative (FN) = 6

From Eq (1) 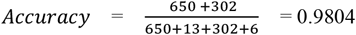

From Eq (2) 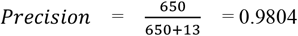

From Eq (3) 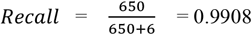

From Eq (4) 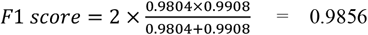

Here Fig. 5 shows that ROC curve along with AUC (Area under curve), AUC shows the difference between positive and negative class, if the AUC is pointing to 1 that means the model performance is good, here our model ROC diagram, the AUC is 1.00 that means it clearly shows that our is model performance is good at differentiating positive and negative classes.

**Fig. 5.**
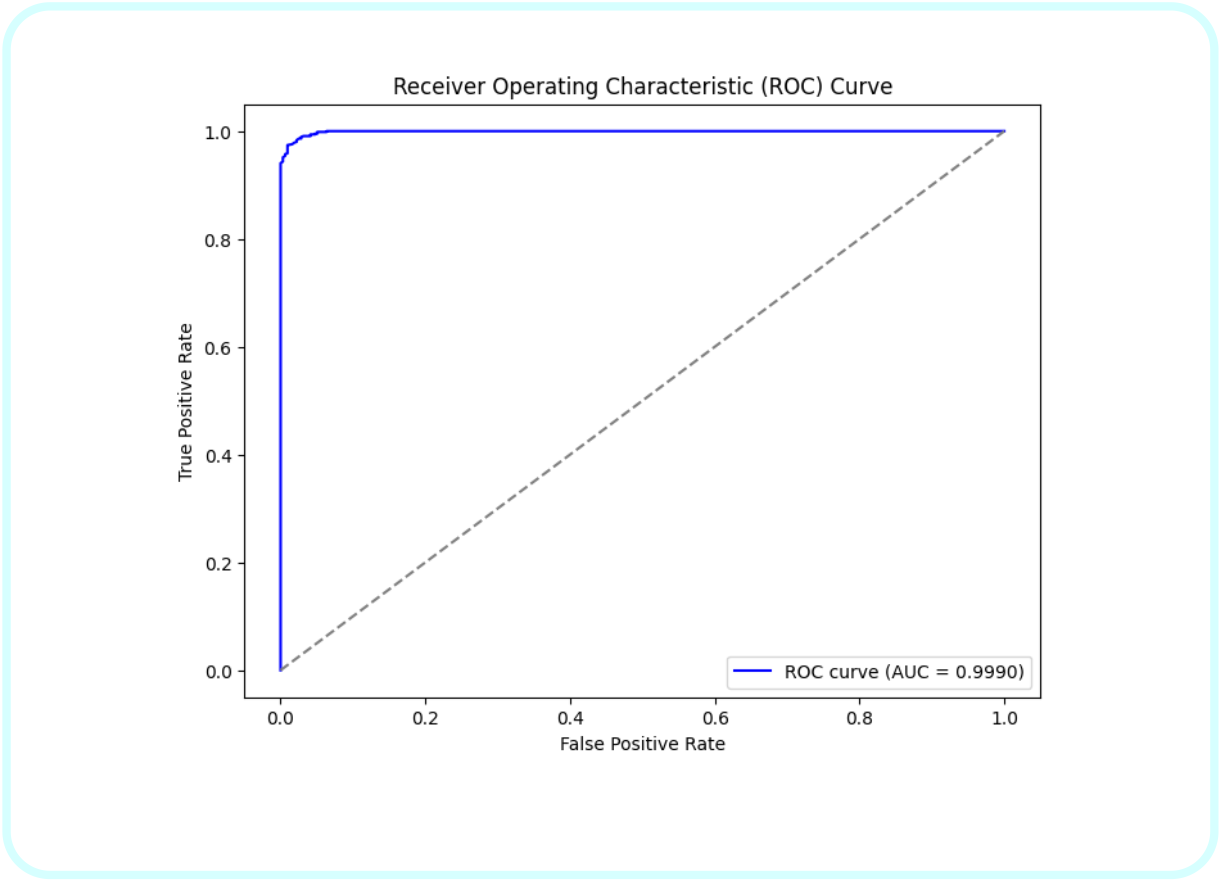
ROC curve for Random Forest Model

Bar Graph: It is a visual comparison of the actual class distribution in the test set and the predicted set, it will help us to diagnose the model’s predictions for any biases.

**Fig. 6.**
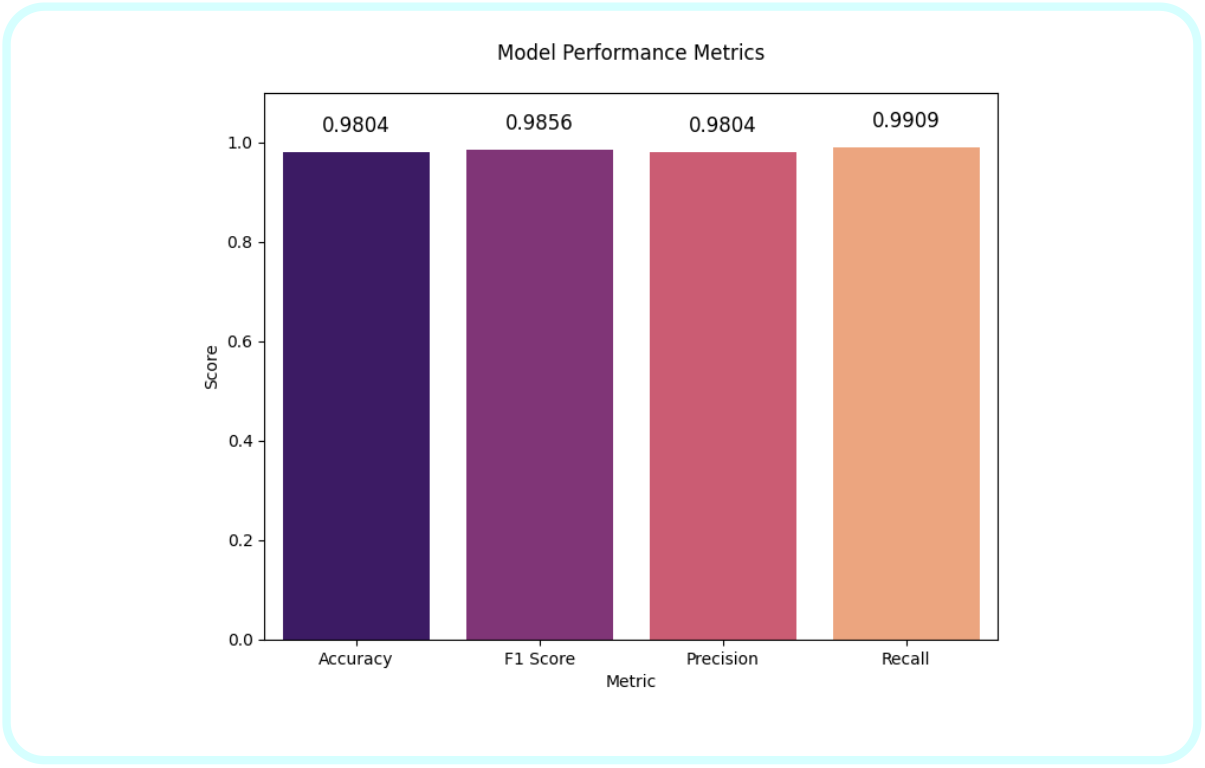
Bar graph for Random forest model

**Table 1.**
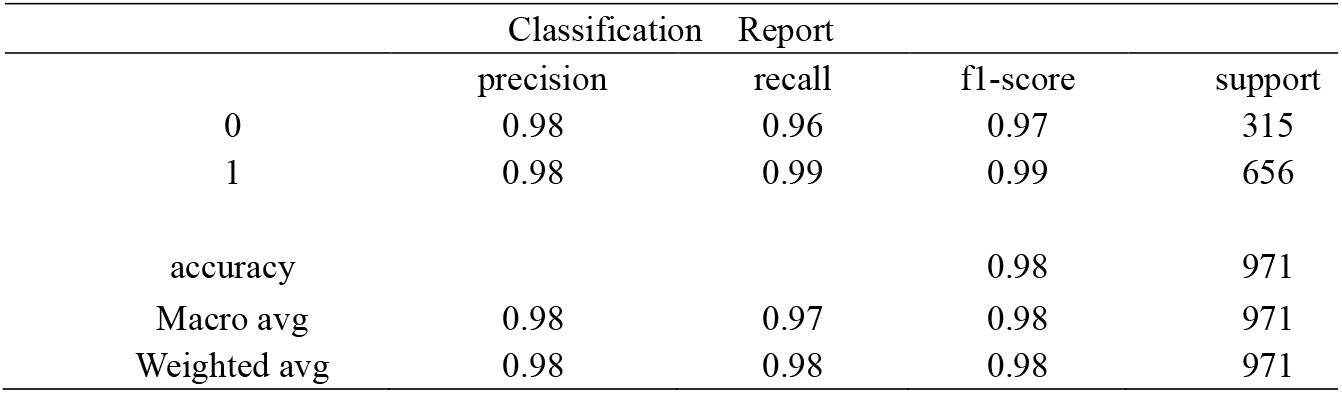
Classification Report for Random forest Model.

The Fig. 7 shows the important feature of word which was evaluate based on the Random Forest model. Here in this diagram, we visualize the top 20 features from the predicted interaction by visualizing we can able to identify and understand the particular terms which are significant during the prediction.

**Fig. 7.**
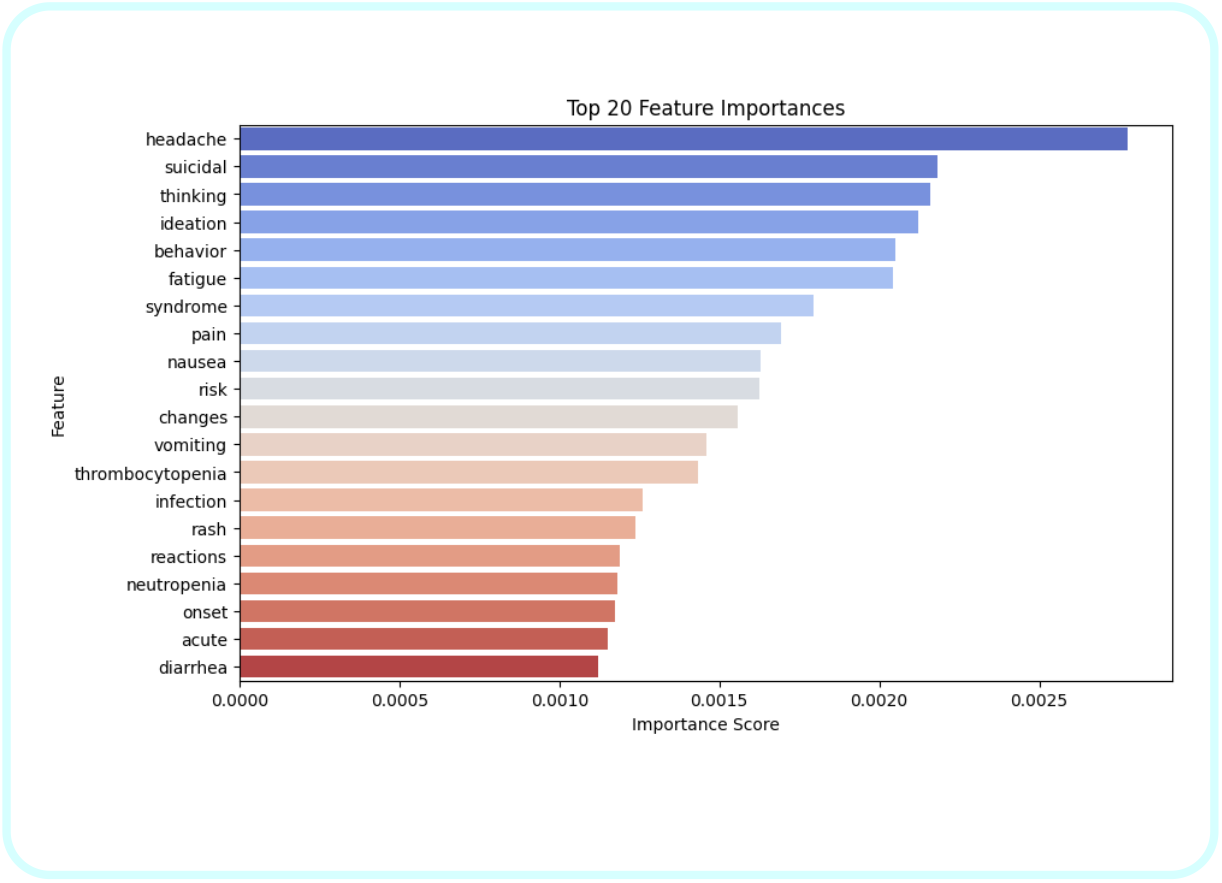
Important Features in Random Forest Model

### 5.2 Gradient Boosting Algorithm and its performance

It is a powerful machine learning technique used mainly for regression and classification tasks [26-30]. It builds a predictive model in a stage wise fashion by sequentially adding models. Boosting models trains models sequentially by new model focusing on residual errors. The initial model makes predictions based on a constant value, such as the mean of the target variable for regression or the most frequent class for classification. Next step is to calculate the residuals based on the difference between the predicted and actual values. These residuals represent the errors. New model is trained based on the residuals of the previous model.The new model is added to the ensemble, and its predictions are combined with previous models.This process is repeated for a set number of iterations until gradually improving the overall model by reducing prediction errors in each step.

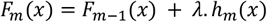

Where,

*F*_*m*_(*x*) - the prediction at m

*F*_*m*− 1_(*x*) - previous model’s prediction.

*h*_*m*_(*x*) - new model trained to predict the residuals.

λ - the learning rate.

Gradient Boosting the disadvantage of overfitting, especially when using complex trees. To prevent this they use regularization techniques like Learning Rate, Tree Depth, Subsampling.

**Fig. 8.**
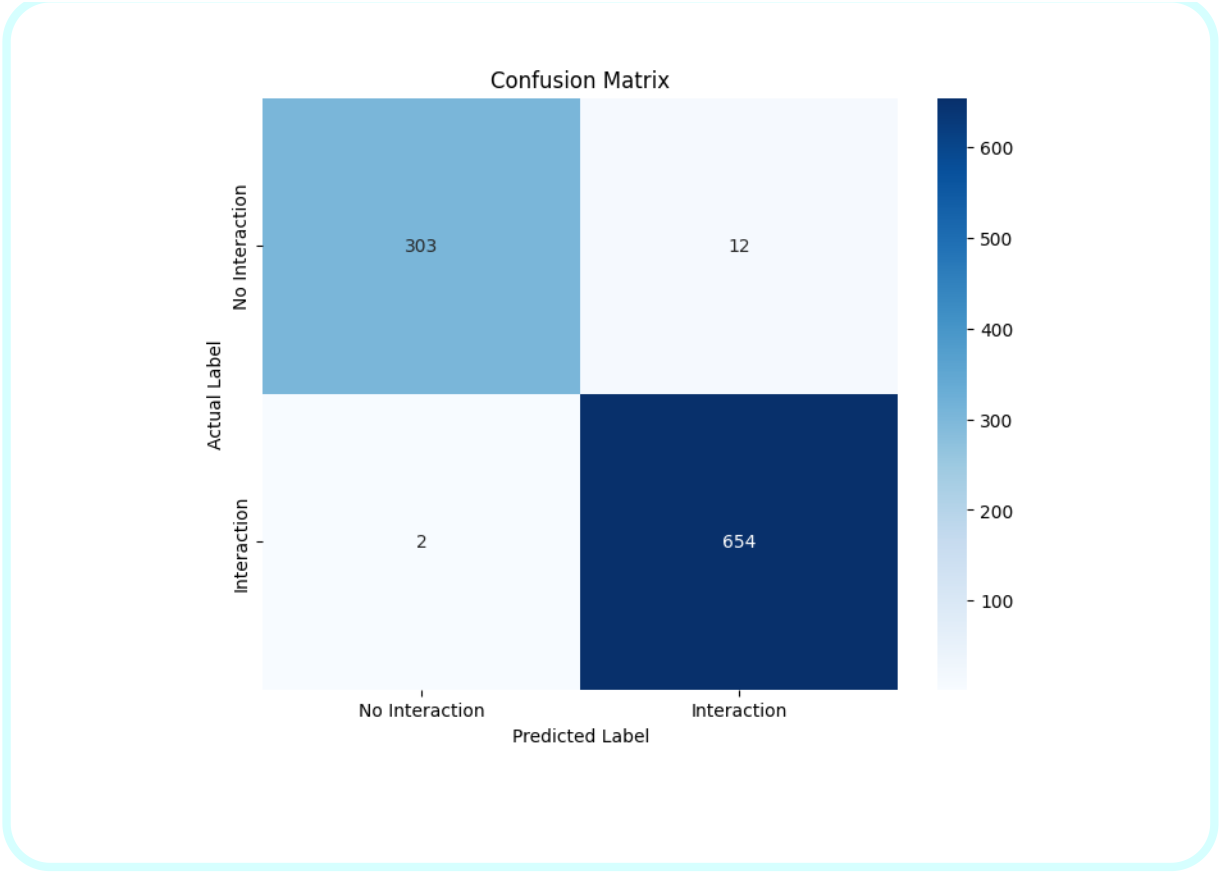
Confusion matrix for Gradient Boosting

True Positive (TP) = 654, False Positive (FP) = 12, True Negative(TN) = 303, False Negative (FN) = 2

Based on the implementation using the respective formulas for accuracy, f1-score,precision and recall we get like:

From Eq (1) Accuracy = 0.9856

From Eq (2) F1-Score = 0.9874

From Eq (3) Precision = 0.9820

From Eq (4) Recall = 0.9970

**Fig. 9.**
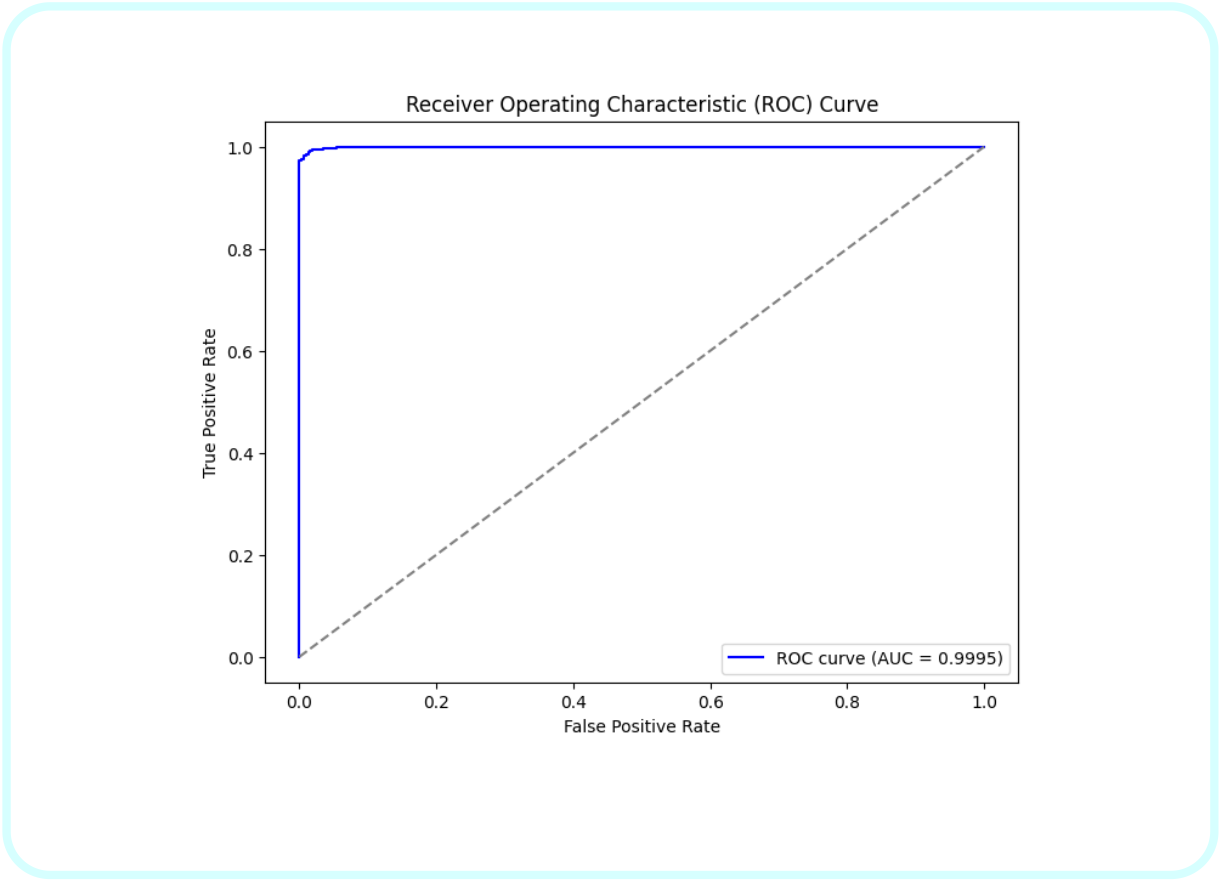
ROC curve for Gradient Boosting

**Fig. 10.**
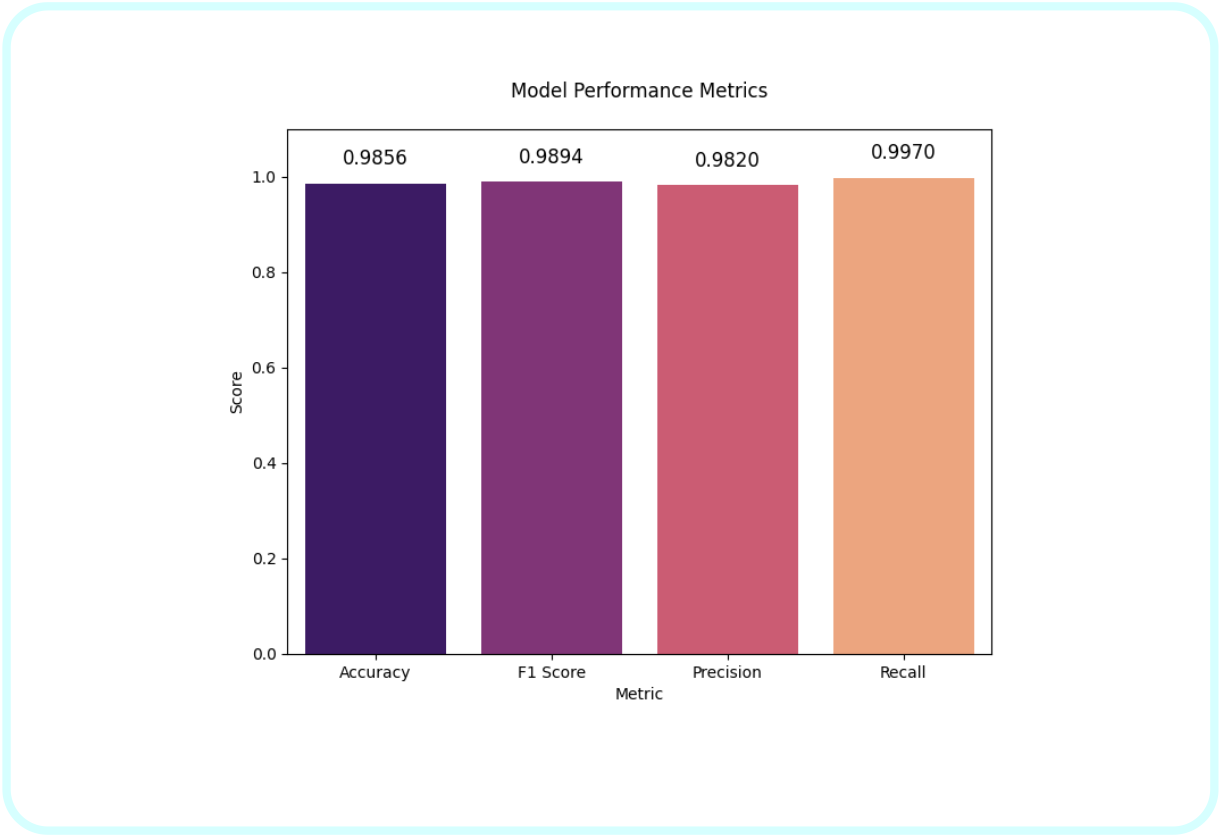
Bar graph for Gradient Boosting

**Fig. 11.**
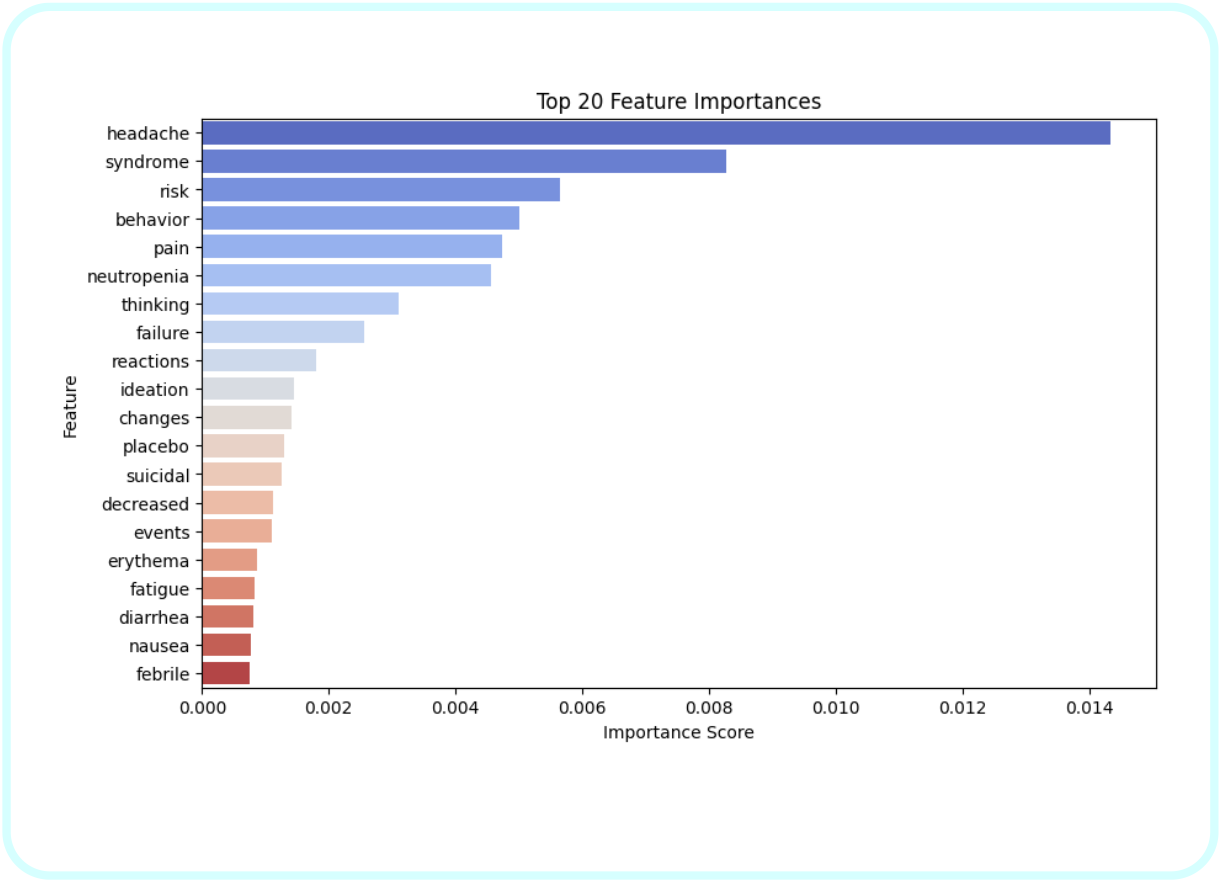
Important Features in Gradient Boosting

**Table 2.** ClassificationReport for Gradient Boosting.

|  |  | Classification Report |  |  |  |
| --- | --- | --- | --- | --- | --- |
|  |  | precision | recall | f1-score | support |
| 0 |  | 0.99 | 0.96 | 0.98 | 315 |
| 1 |  | 0.98 | 1.00 | 0.99 | 656 |
| accuracy |  |  |  | 0.99 | 971 |
| Macro avg |  | 0.99 | 0.98 | 0.98 | 971 |
| Weighted avg |  | 0.99 | 0.99 | 0.99 | 971 |

### 5.3 Support Vector Machine (SVM) and its performance

It is a powerful and versatile method of supervised machine learning algorithm especially used for classification and regression tasks [31-37]. It works by finding the hyperplane that best divides the dataset into different classes. It has key concepts such as hyperplane, support vectors, Margin, Linear SVM, Non-linear SVM (Kernel Trick), Soft Margin and Regularization. Consider a binary classification problem with two classes (+1 and −1). The SVM algorithm would find a hyperplane that maximizes the margin between the two classes. If the data is not linearly separable,slack variables are introduced to allow margin violations.

**Fig. 12.**
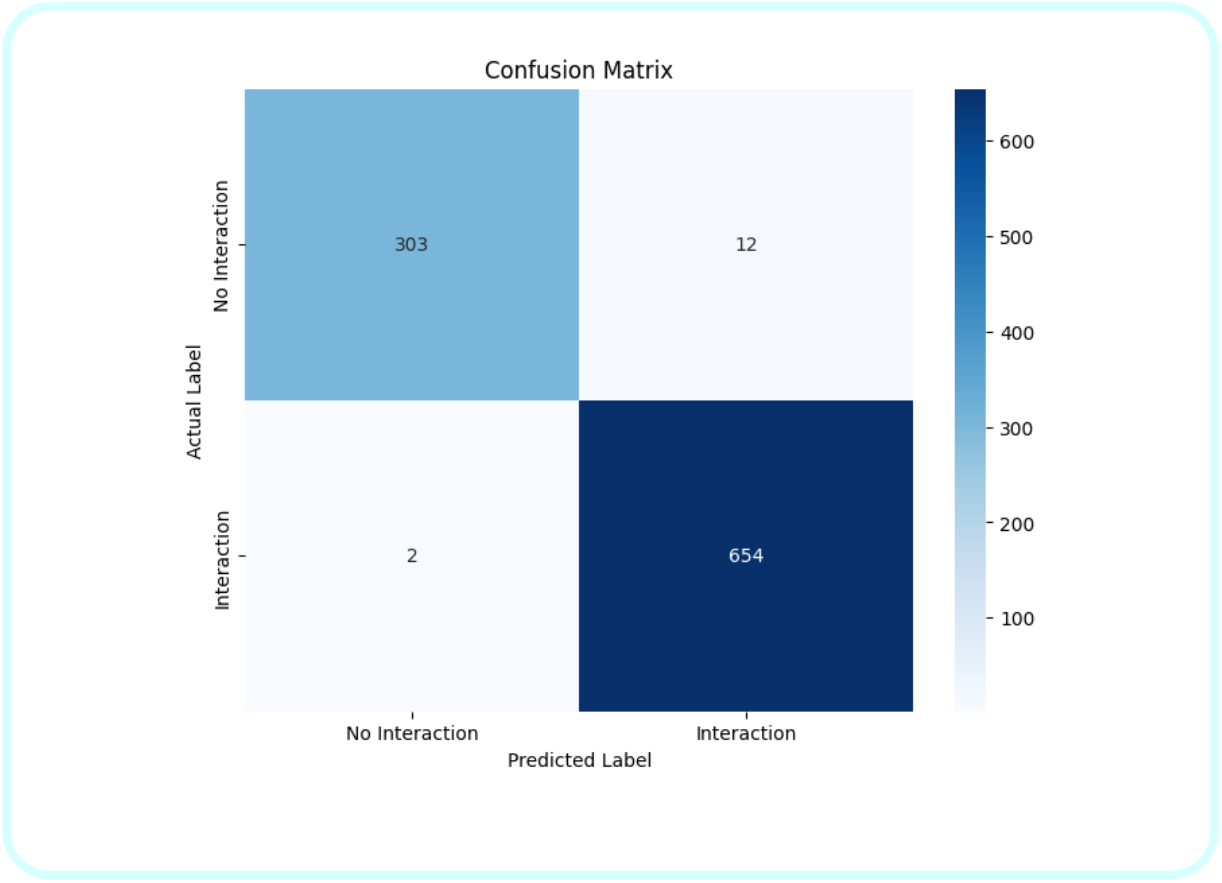
Confusion Matrix for SVM

**Fig. 13.**
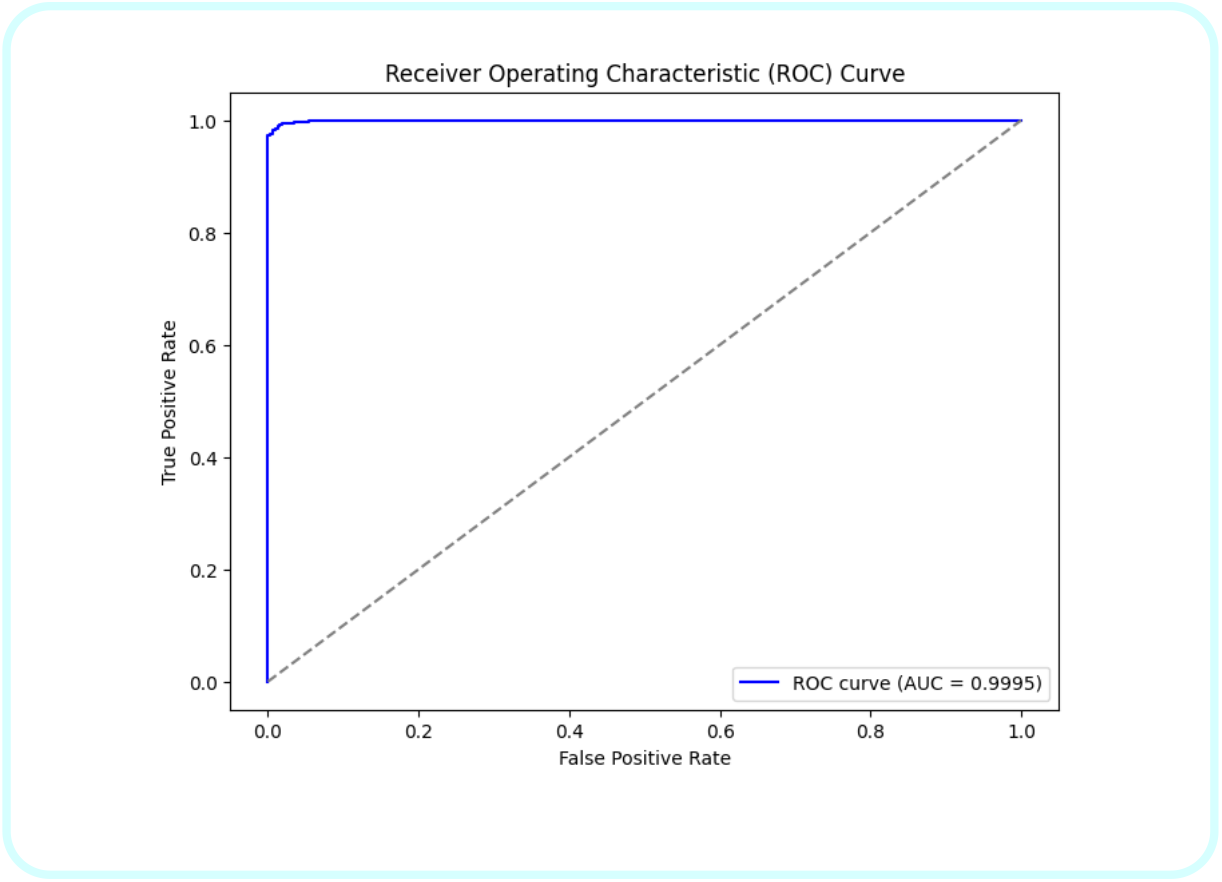
ROC curve for SVM

True Positive (TP) = 654, False Positive (FP) = 10, True Negative(TN) = 305, False Negative (FN) = 2 Based on the implementation using the respective formulas for accuracy, f1-score,precision and recall we get like:

From Eq (1) Accuracy = 0.9876

From Eq (2) F1-Score = 0.9909

From Eq (3) Precision = 0.9849

From Eq (4) Recall = 0.9970

**Fig. 14.**
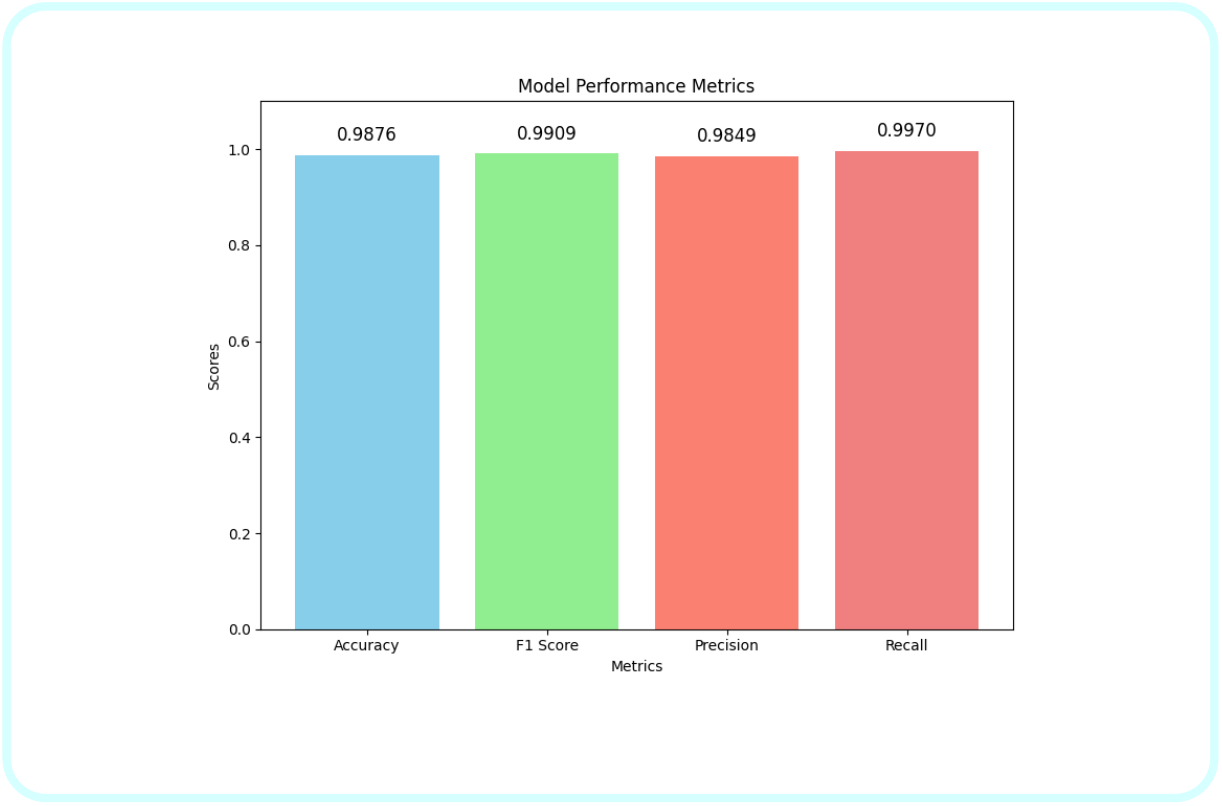
Bar Graph for SVM

**Table 3.**
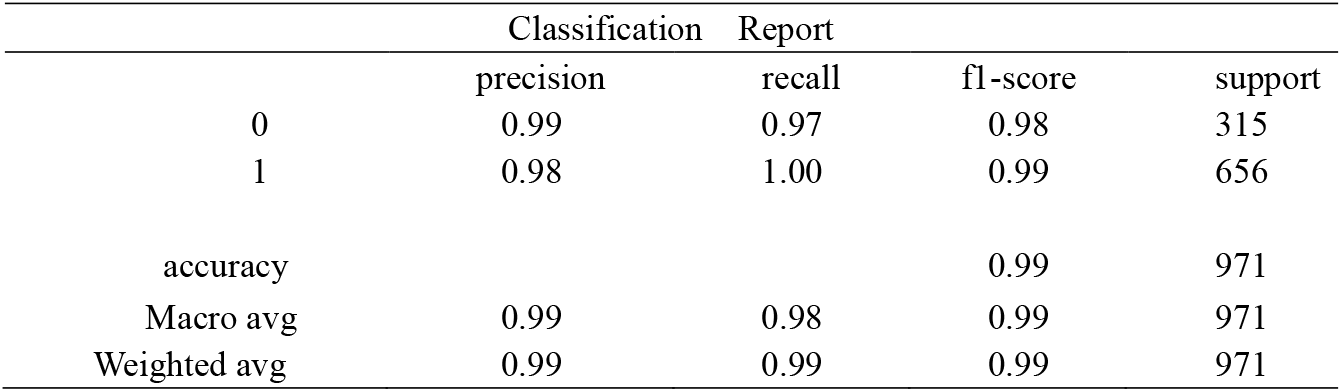
Classification Report for SVM.

|  | Classification |  | Report |  |
| --- | --- | --- | --- | --- |
|  | precision | recall | f1-score | support |
| 0 | 0.99 | 0.97 | 0.98 | 315 |
| 1 | 0.98 | 1.00 | 0.99 | 656 |
| accuracy |  |  | 0.99 | 971 |
| Macro avg | 0.99 | 0.98 | 0.99 | 971 |
| Weighted avg | 0.99 | 0.99 | 0.99 | 971 |

### 5.4 BioBERT and its performance

BioBERT (Bidirectional Encoder Representations from Transformers for Biomedical Text Mining) is a pre-trained language model particularly focused on biomedical domain [38-40]. BERT (Bidirectional Encoder Representations from Transformers), one of the most important models in natural language processing (NLP), because it is specifically designed to handle biomedical and clinical literature. BERT outperform other traditional models by processing words in all direction. BioBERT is designed to handle biomedical text like Named Entity Recognition (NER), Relation Extraction, Questions related to biomedical. BioBERT retains the transformer architecture model from BERT. It consists of layers of self-attention mechanisms that allow it to process each word simultaneously attending every other word. The model’s parameters include number of transformer layers, heads, and hidden units are identical to those of BERT, but the difference is pre-training corpus. In our proposed model for BioBERT, the BioBERT embeddings represent the side effects of drug pairs, these drug pairs are then as concatenated embeddings. A simple neural network model with one hidden layer was trained to classify whether the drug pairs have common side effects.

**Fig. 15.**
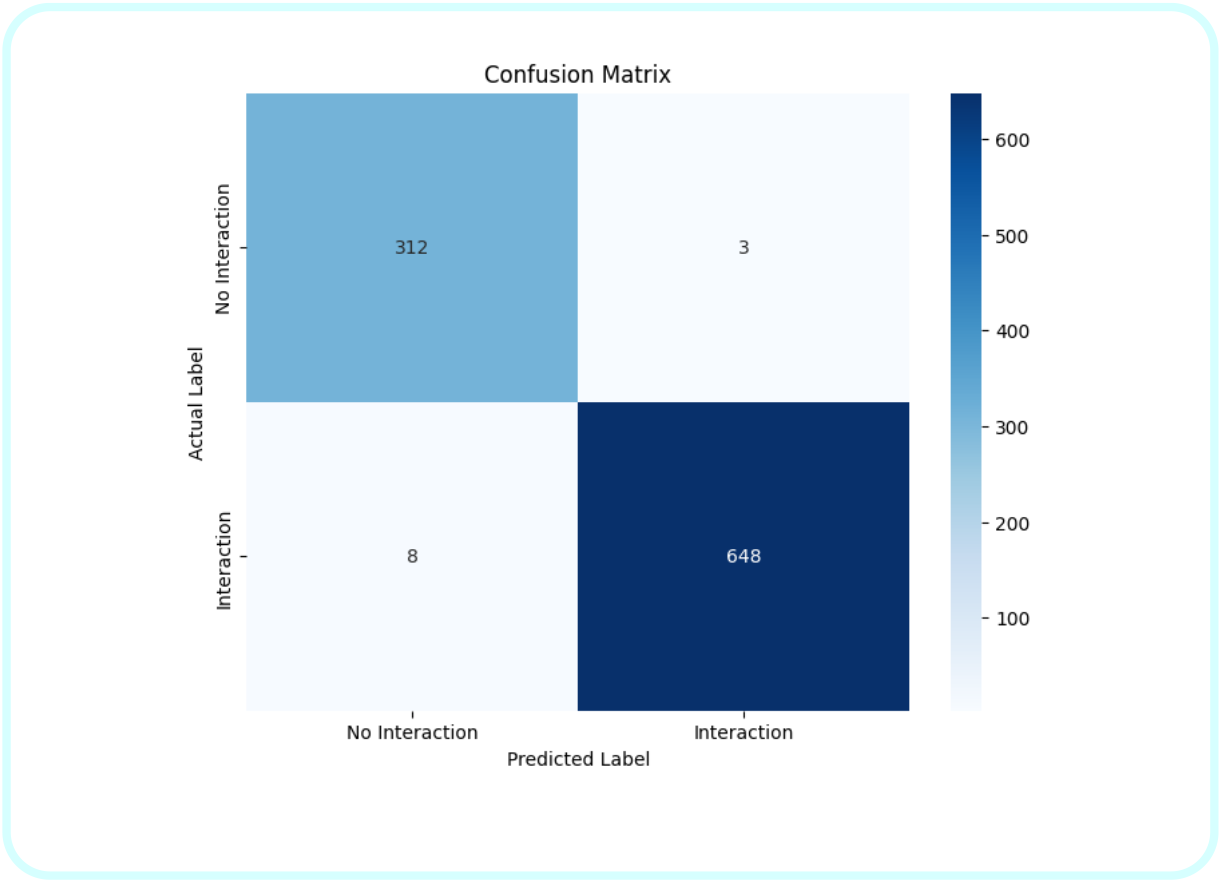
Confusion Matrix for BioBERT

**Fig. 16.**
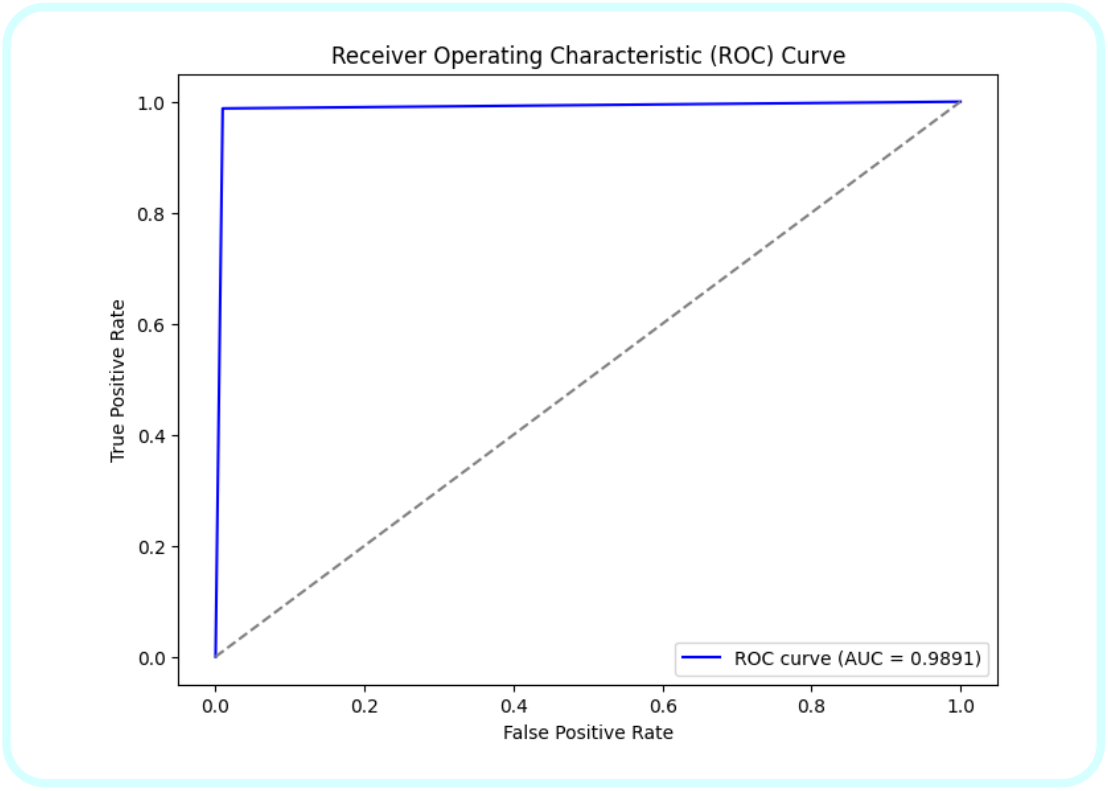
ROC curve for BioBERT

True Positive (TP) = 648, False Positive (FP) = 3, True Negative(TN) = 312, False Negative (FN) = 8

Based on the implementation using the respective formulas for accuracy, f1-score,precision and recall we get like:

From Eq (1) Accuracy = 0.9887

From Eq (2) F1-Score = 0.9916

From Eq (3) Precision = 0.9954

From Eq (4) Recall = 0.9878

**Fig. 17.**
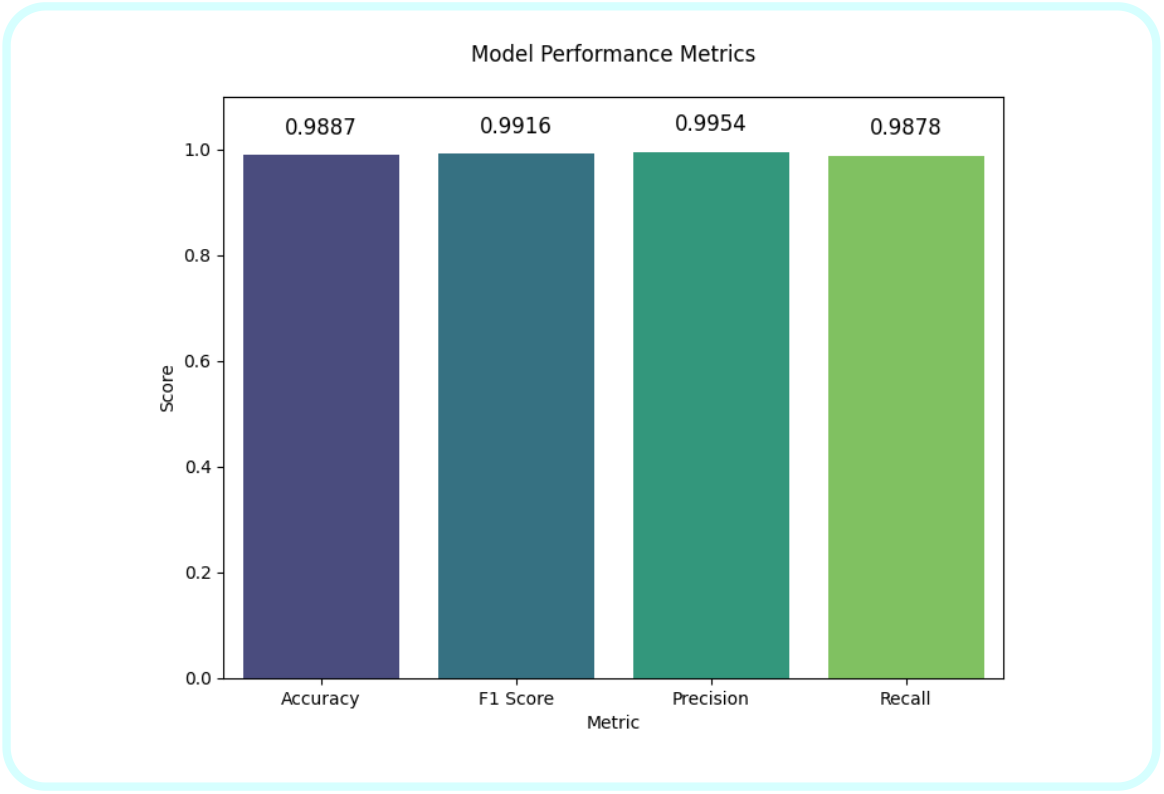
Bar graph for BioBERT

**Table 4.**
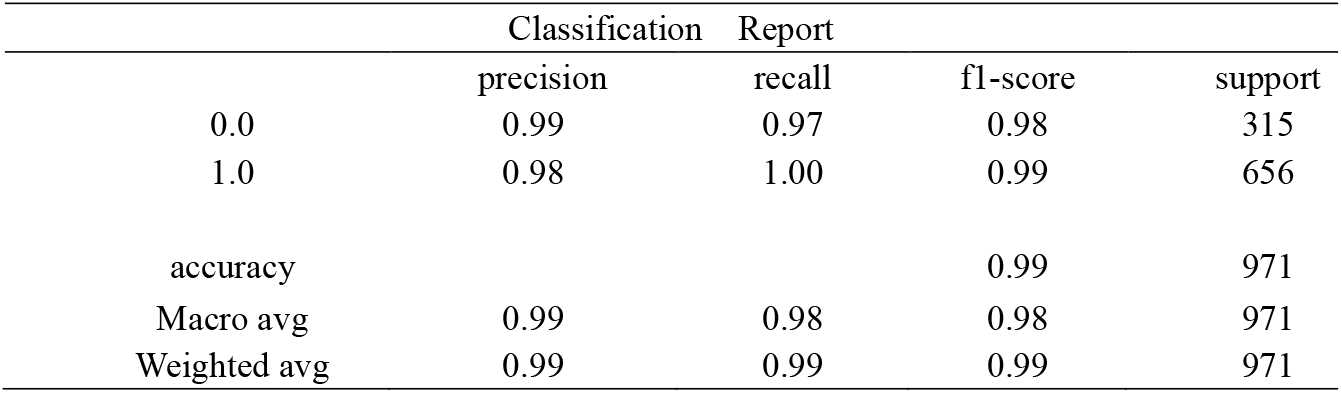
Classification Report for BioBERT.

|  | Classification |  | Report |  |
| --- | --- | --- | --- | --- |
|  | precision | recall | f1-score | support |
| 0.0 | 0.99 | 0.97 | 0.98 | 315 |
| 1.0 | 0.98 | 1.00 | 0.99 | 656 |
| accuracy |  |  | 0.99 | 971 |
| Macro avg | 0.99 | 0.98 | 0.98 | 971 |
| Weighted avg | 0.99 | 0.99 | 0.99 | 971 |

## VI. Final Comparison

This table show the evaluation metrics we achieved from training the various model for DDI Extraction.

**Table 5.**
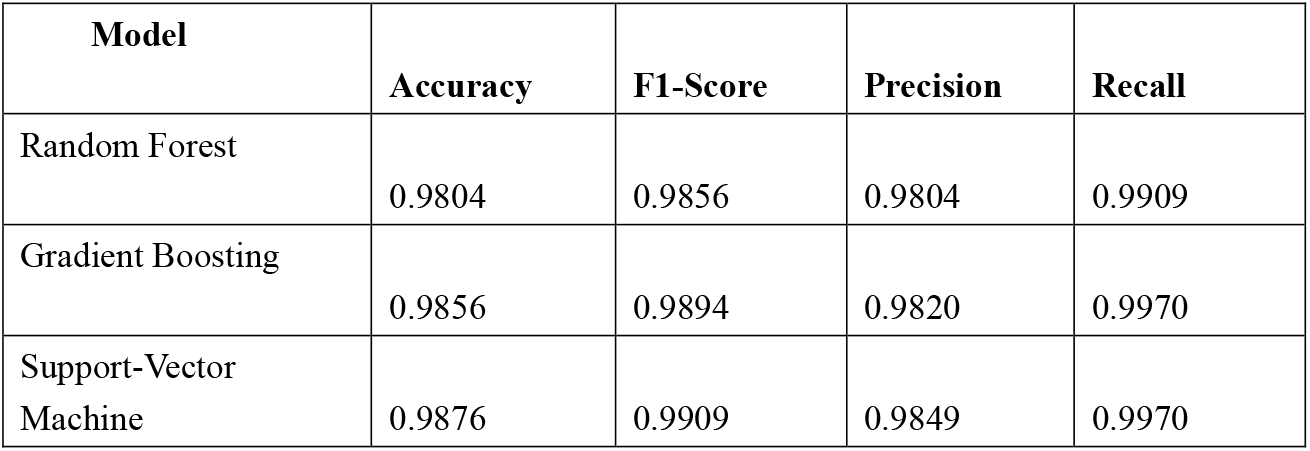
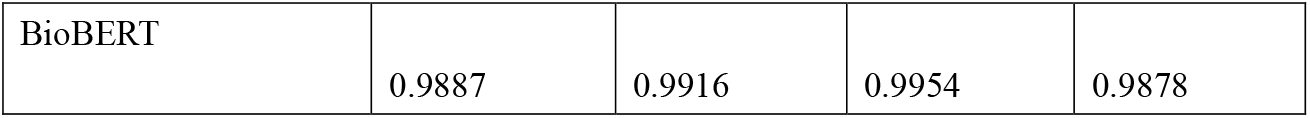
Final Comparison Table.

## VII. Discussion

Our model is designed to classify drug-drug interactions using drugs and their Adverse reaction from the TAC 2017 dataset, we used TF-IDF for text processing, After TF-IDF it fed into normalization for balancing the unique features from the data for further processing, we proposed three machine learning models and one transformer model namely Random Forest, Gradient Boosting, Support vector machine and BioBERT for DDI (Drug-Drug Interaction) extraction. However our proposed model is aimed to predict and classify drug interaction accurately, along with increasing training and Visualizing evaluation through metrics to ensure that our model can able to understand and improve the performance. The final predicted model gives the side effects of combining drug A and drug B. For Example, combining gattex and fycompa may lead to the following side effects: pyrexia, reactions, fatal, oral, hypotension.

## VIII. Conclusion and Limitation

### 8.1 Conclusion

In this study we proposed Multiple approach for the classification for drug and their adverse reaction by using the TAC 2017 ADR Dataset. We designed three machine learning model with TF-IDF (Term Frequency-Inverse Document Frequency) and one transformer model for classification namely Random Forest, Gradient Boosting, Support Vector Machine and BioBERT(Bidirectional Encoder Representations from Transformers for Biomedical Text). For Random Forest we achieved an Accuracy of 0.9804, an F1-Score of 0.9856, a precision of 0.9804, a recall of 0.9909, for Gradient Boosting we achieved an Accuracy of 0.9856, an F1-Score of 0.9894, a precision of 0.9820, a recall of 0.9970, for Support Vector Machine we achieved an Accuracy of 0.9876, an F1-Score of 0.9909, a precision of 0.9849, a recall of 0.9970, and for BioBERT we achieved an Accuracy of 0.9887, F1-score of 0.9916, a precision of 0.9954, a recall of 0.9878. From this observation BioBERT model outperforms compared to the other Machine learning models with few increasing number of percentages with visualization metrics. The experimental results shows that our proposed model yields better performance for the classification for drug and their adverse reaction.

### 8.2 Limitation

In this study, we done the classification for drug and their adverse reaction using four models, even though they yield promising accuracy for the classification for DDI(Drug-Drug Interaction) extraction but it is not considered as a perfect model. In future, we will try to implement a new approach with new dataset which comprises chemical molecule along with DDI information for giving a new insights for health care.

## Acknowledgements

We would like to express our sincere gratitude to the professors, their guidance and support gave valuable insights throughout this research. We would also like to appreciate the professionals and organizations who provided valuable insights and data in the areas of the TAC 2017 dataset, Machine learning models, and Transformer models for Drug-Drug Interaction. Their hard work gives us clear insights into our findings and classification. A sincere thanks to our colleagues who are always active in the research process. We will take responsibility if there are any mistakes in the document.

